# Hypoxia Imaging as a Radiomics Signature in Tumors Using Novel ^19^F-Eu-Based Contrast Agents for Magnetic Resonance Imaging

**DOI:** 10.1101/2025.10.11.681845

**Authors:** Susan A. White, S. A. Amali S. Subasinghe, Jonathan Romero, Md. Abul Hassan Samee, Jason T. Yustein, Robia G. Pautler, Matthew J. Allen

## Abstract

The ability to accurately identify and characterize regions of hypoxia has been an active area of study due to the biological ramifications of low oxygen tension in different pathological conditions, including inflammation, infections, wound healing, cardiovascular conditions, kidney and pulmonary diseases, hepatic and neurological toxicities, and cancer. Although hypoxia contributes a significant role in these conditions, the ability to accurately and instantaneously monitor the presence of hypoxia *in vivo* and correlate comprehensive analysis of hypoxia-dependent molecular signatures in response to treatments regimens is lacking. With the advent of hypoxia-responsive contrast agents for magnetic resonance imaging (MRI), including the recent development and characterization by our team of a novel probe that involves both ^19^F and Eu^II^, the capability to integrate direct hypoxia imaging in real-time with new technologies that enable spatial transcriptomic profiling has become a possibility. To assess the capability of this agent for studying hypoxia, we used osteosarcoma as a model. In this preliminary study we demonstrate two major results: First, we show that convection-enhanced delivery (CED) is a reliable and robust methodology to distribute MRI contrast agents throughout a tumor. Second, we show that integration of direct hypoxia-detecting imaging modalities and spatial biology enables real-time *in vivo* insights into biology and identification of biomarkers directly applicable to disease development and response to therapy. The ability to identify and define hypoxia-mediated biology using direct rather than indirect MRI methods has extremely significant implications in the care of a wide-range of pathological conditions. Consequently, the framework outlined in these preliminary studies is applicable to other pathological conditions and provides the basis for direct *in vivo* hypoxia detection, monitoring, and biological analyses with translational applications to patient care and management.

## Introduction

Hypoxia, or low oxygen tension, contributes to different pathological conditions, including inflammation, infections, wound healing, cardiovascular conditions, kidney and pulmonary diseases, hepatic and neurological toxicities, and cancer.^1–26^ For example, intratumoral hypoxia disrupts the homeostasis of reduction and oxidation (redox), altering tumor metabolism and downstream molecular effectors that contribute to tumor development and progression.^1–41^ These changes influence tumor aggressiveness, invasiveness, metastasis, and resistance to treatment, making hypoxia a significant target for biomedical imaging and radiomics.^2–7,10–31,38,39,42–47^

The accurate, noninvasive imaging of hypoxia is critically important because hypoxia and its consequences are attractive targets of several new drug therapies,^2–7,10–20,24–27,36,37,44,48–50^ and evaluation of the effectiveness of these therapies and their mechanism of action, even at the preclinical stage, would be enhanced with the ability to directly and instantaneously image hypoxia. Consequently, various *in vivo* redox-responsive techniques have been investigated to image hypoxia: Positron emission tomography is in current clinical and preclinical use,^3,7,14,26,43,44,51–58^ and although, positron emission tomography can map hypoxia, this technique offers no anatomical details on its own, thus requiring a second imaging modality. Also, it often involves probes with poor pharmacokinetics, and it involves ionizing radiation, limiting its use in repeated imaging, especially with pediatric patients.^46,55–61^ Electron paramagnetic resonance imaging provides direct measurement of tissue oxygen levels, but the number of these scanners is extremely limited and the technique offers no anatomical details on its own, requiring separate sequential imaging with a second modality.^14,53,58–62^ Perfluorocarbon emulsion oximetry agents for ^19^F-MRI and oximetry agents for ^1^H-MRI have low aqueous solubility that restricts biodistribution to areas orthogonal to water-soluble molecules, and ^1^H-NMR oximetry agents often overlap with fat signals, complicating the interpretation of data.^14,53,55,60,61,63–65^ Additionally, hypoxia can be visualized with *ex vivo* immunohistochemistry; however, this approach is invasive in that it requires biopsy and without a way to image the entire tumor, can be misleading due to tumor hypoxia heterogeniety.^14,26,47,51,53,59,61,66,67^ Redox-responsive contrast agents for MRI with ligand-, organic-radical-, or transition-metal-based contrast agents have been used for *in vivo* imaging,^68– 85^ but these agents produce contrast enhancement in both hypoxic and normoxic regions, never achieving a truly silent off-state. Indirect MRI methods have been evaluated to determine their efficacy in monitoring *in vivo* hypoxia. Diffusion-weighted imaging alone has been shown to not correlate with hypoxia;^86–93^ however, dynamic contrast enhancement (DCE) that measures the perfusion of an extracellular MRI contrast agent into a tissue has had some success measuring hypoxia.^86,87,94–97^ However, because DCE is based on vascular flow as an indirect reporter of hypoxia, it can only detect hypoxia due to vascular changes^98^ and also inherently cannot distinguish hypoxia from hypoxemia. Hypoxia is a reduction in levels of tissue oxygenation, whereas hypoxemia is a decrease in the partial pressure of blood oxygen. The most common mechanism of hypoxemia is a mismatch of ventilation and perfusion. This inability to differentiate hypoxemia from hypoxia is critical because as in the case of cancer, hypoxia is oftentimes observed as an indicator of malignancy, but chemotherapy treatments have been noted to induce hypoxemia.^99–104^ Because DCE cannot readily distinguish between hypoxia and hypoxemia, the hypoxia readouts generated from DCE may be misleading and negatively impact patient care, especially during chemotherapy treatment. Thus, creating technologies that directly measure hypoxia, and are not perfusion based, are critically needed. Although each reported approach has enhanced the understanding of redox biology, the limitations of these methods fuel gaps in knowledge that are addressed by using a ^19^F-Eu-based contrast agent.

To perform well as a redox-active probe for hypoxia, a complex must have a molecular switch that can dynamically alter contrast enhancement at detectable levels in response to low-oxygen conditions. However, the clinically used ion Gd^III^ is locked into a single oxidation state under physiological conditions and, therefore, is unable to act as a metal-based redox switch to detect hypoxia. The +2 oxidation state of europium (Eu^II^) is isoelectronic (seven unpaired electrons) with Gd^III^, and the two ions have similar relaxivities that characterize the ability to enhance contrast in MRI.^105–115^ The similar relaxivities cause Eu^II^-based contrast agents to be used in comparable concentrations to Gd^III^-based contrast agents for MRI.^106^ Additionally, Eu^II^ irreversibly oxidizes to Eu^III^ under physiological conditions, and Eu^III^ is completely *T*_1_-silent at imaging-relevant concentrations:^1,110,112,116,117^ The irreversible nature of the couple avoids confusion from dynamic processes. Eu^II^-containing complexes respond to hypoxia in MRI directly through the oxidation states of the metal ion, whereas Gd^III^-based redox systems involve redox chemistry of ligands, causing some level of contrast enhancement from Gd^III^ regardless of the activity of the ligand.^73^ For non-Eu^II^ metal-based systems, like Fe^II/III^ and Mn^II/III^, the off-state is not *T*_1_-silent but only provides a lower degree of positive contrast enhancement that is higher than the off-state of negative controls in phantom imaging.^68–80,82–84,118–122^ Therefore, the metal-centered response together with the unique electronic properties of Eu^II^ and Eu^III^ cause redox-active Eu-based contrast agents for MRI to overcome critical limitations of Gd^III^-, transition-metal-(for example Mn or Fe), and ligand-based systems. Although the use of Eu^II^ for oxygen sensing in MRI was proposed at the turn of the century,^105,106^ Eu^II^ was considered incompatible with living systems until recent studies demonstrated the ability of Eu^II^ to enhance contrast as a function of oxygen content *in vivo*.^1,116,123^ Subsequently, we imaged hypoxia in preclinical models of osteosarcoma,^124^ and our *in vivo* results in healthy and tumor-bearing mice demonstrate that Eu^II^-based systems have the potential to address important gaps for imaging hypoxia.

As therapies for the treatment of cancer become more diverse, one of the greatest challenges is the optimization of delivery systems to tumors. The current development of novel gene therapies, the ongoing development of degradomers as well as other novel therapeutics make the implementation of delivery technologies for such therapeutics extremely important. One delivery solution that has received recent attention is that of convection enhanced deliver (CED).^125–130^ This technique enables repeated and targeted delivery of agents to tumors through an implanted cannula thereby circumventing systemic effects. CED is an innovative methodology by which a catheter is placed into a tumor to infuse agents.^131–133^ CED uses an infusion pump to create a positive pressure gradient by which convective transport of the infused agent is established. The advantages of CED include producing a well-targeted and high local concentration of agent while limiting systemic toxicities. CED is superior to direct tumor injections that rely solely upon diffusion.^131–133^ Direct tumor injection has a limited volume of distribution whereas CED can cover a much broader area depending upon infusion times and rate. An additional limitation of direct tumor injection includes undesirable systemic delivery of agent and potential off target effects, which are not issues when using CED. There have been multiple successful clinical trials using CED in cancer patients that include infusion of chemotherapeutic agents, viral vectors, and monoclonal antibodies.^134–140^ Moreover, CED is already an FDA-approved technique for use in patients.^141–150^ The advantages of CED over direct or even systemic injections make CED the ideal choice for administration of ^19^F-Eu-based imaging agents. Further the combination of ^1^H and ^19^F signal enables ratiometric imaging that accounts for diffusion in the proposed studies. Indeed, because CED is making its way into the clinic for the local delivery of therapeutics to tumors, this delivery system can also be used for the local delivery of imaging agents.

In the studies described here, we first validated the use of convection enhanced delivery (CED) to distribute MRI contrast agents throughout tumors. Then, we used CED to infuse a mouse model of the previously established hypoxic osteosarcoma with our ^19^F-Eu-based agent. Finally, to validate that the agent was aligning with regions of hypoxia, we then used Nanostring spatial transcriptomic technology to confirm regions of hypoxia versus normoxia. Using statistical association, we were able to confirm the correlation of gene clustering between hypoxic and normoxic regions of interest (ROI) identified by our novel ^19^F-Eu-based agent and differential gene expression between hypoxic and normoxic ROI. These data in this preliminary study confirm the utility of using CED to distribute MRI contrast agents throughout tumors. Additionally, our data illustrate the potential utility of these novel ^19^F-Eu-based contrast agents for MRI in denoting hypoxia in tumors. These results demonstrate that CED MR imaging with ^19^F-Eu-based contrast agents can be applied as candidate biomarkers for tumor phenotypes, including responsiveness to therapeutic regimens.

## METHODS

### Intratibial Orthotopic Syngeneic Model, Tumor Monitoring, and Randomization

mCherry-labeled murine osteosarcoma cells (5 × 10^5^ cells of F420, F408, or F331) were injected intratibially into C57BL/6 mice. Tumors were monitored and volumes measured biweekly. When primary tumors reached 0.25–0.5 cm^3^ [using *V* = *A* × *B*^2^/2 *(A = largest diameter; B = smallest diameter*)] were used for this preliminary study.

### Using three-dimensional printing of tumor molds

From the three-dimensional MRI dataset of each tumor, we used Osirix to segment the tumors. From the DICOM files, we used Fusion 360 software to generate.stl files. We then used the Ultimaker Cura software to convert.stl files into printable molds with our high-resolution 3D printer (Ultimaker S3). Molds included slots to enable removing tissue sections that align with MRI for Nanostring Digital Spatial Profiling.

### Validation of imaging with Pimonidazole staining (hypoxyprobe)

To validate hypoxia imaging, we performed pimonidazole (sold as Hypoxyprobe-1) staining for *in situ* detection of hypoxic regions within the resected sarcoma specimens. Specifically, pimonidazole (60 mg/kg per manufacture recommendation) was injected via tail vein 60–90 min prior to sacrifice of the mouse.

*CED of the* ^*19*^*F-Eu-based probe:* For CED, we used a Harvard Apparatus infusion system, to infuse 120 μL at rate of 1.5 μL/min. An infusion catheter was placed so the catheter tip was in the center of the tumor.

### Hypoxia MRI

All preclinical imaging was conducted on a 9.4 T Bruker AV NEO, 20 cm bore MRI with a dual tuned Bruker ^19^F/^1^H 40 mm volume coil. Mice were induced with 5% isofluorane and transferred supine to the animal-imaging holder, and then transferred to the imaging instrument where 2% isofluorane was administered continuously via nose cone. A pressure-sensitive pillow was placed on the abdomen to monitor respiration. Temperature was monitored using a rectal probe and maintained at 37 °C using a heating system. For assessment of hypoxia after CED, we generated tumor *T*_1_ maps and measured ^19^F signal using chemical shift imaging one-hour post CED. A single-pulse ^19^F spectrum was acquired to monitor the presence of detectable ^19^F. *T*_*1*_ *and T*_*2*_ *measurements*. Standard rapid acquisition with relaxation pulse sequences were used, and *T*_1_ and *T*_2_ maps (with variable repetition times and echo times) were acquired and processed using Paravision 360. ^*19*^*F imaging (chemical shift imaging)*. ^19^F-MR spectroscopic imaging was performed using a fast spin echo imaging sequence with the following imaging parameters: field of view = 32 mm × 32 mm; image size = 16 × 16 voxels; slice thickness = 15 mm; repetition time = 250 ms; echo time = 0.647 ms; number of averages = 8; 1024 FID data points, spectral width = 25 kHz.

### Preprocessing and quality control of Nanostring Digital Spatial Profiler data

Initial processing and quality control of the data was performed following the Nanostring standard pipeline.^151–153^ Recent studies validated the robustness of this technology in quantifying protein or transcript abundance from selected ROI.^151,153^ Data was inspected to ensure high levels (as prescribed by Nanostring and recent literature)^154^ of aligned read count, sequencing saturation, and number of detected genes above the limit of quantitation, compared to negative probes from Nanostring. Next, as recommended in a recent benchmarking of Nanostring data normalization methods, the RUVSeq method was applied to normalize the data and remove potential technical variabilities.^155,156^ *Preparation of formalin-fixed, paraffin-embedded tumors:* Murine osteosarcoma tumors were fixed in neutral buffered formalin and PhosStop to preserve phosphorylated proteins. Tumor samples were then paraffin embedded with standard protocols of dehydration, clearing, and embedding.

### Nanostring Data

Eight ROI spanning conditions from normoxic to hypoxic were selected. Within each region, cells were stained for smooth muscle actin (SMA), a marker for smooth muscle differentiation in tumors, and CD31, a marker for endothelial cells. Three distinct whole-transcriptome expression profiles were generated per region by segregating cells into SMA-positive, CD31-positive, and SMA-negative/CD31-negative populations. In effect, each region was subdivided into three spatial regions, each represented by its own transcriptome profile. For simplicity, we refer to each of these transcriptome profiles as a distinct region of interest.

### Clustering of ROI

t-SNE plots were produced, and clustering of ROI was performed using Nanostring’s proprietary data analysis tool. The resulting visualizations were then exported.

### Gene clustering

We used hierarchical clustering taking the distance between rows (or columns) *x* and *y* being: 1 − pearson(*x, y*), where pearson(*x, y*) denotes the Pearson correlation coefficients between *x* and *y*.

### Differential gene expression testing

A two-tailed Welch’s t-test was conducted between hypoxic and Normoxic ROI. The test accommodated the possibility of unequal variances between groups. *Proportions of cell types in ROI using single-cell signatures:* For each region of interest *R* and cell type *C*, we first computed an abundance score *α*_*C,R*_, which is the average of Z-scores of the marker genes of *C* in *R*. This score *α*_*C,R*_ is then converted to a proportion *f*_*C,R*_ as: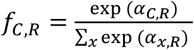.

## RESULTS

First, we tested CED with the Gd^III^-based MRI contrast agent Magnevist and determined that through this delivery method, we can infuse contrast agent throughout the entire tumor rather than just portions of the tumor as would occur through intratumoral injection (Figure 1). Next, we injected the ^19^F-Eu-based probe (Figure 2) into a separate tumor using CED. For that tumor, we combined two published methods to use a high-resolution 3D printer to create a mold that enables sectioning of tissues to match MRI datasets (Figure 3).^157,158^ We also determined that using the software, RadPathFusion,^158,159^ would ensure that tissue sections aligned with MRI images (Figure 3). We used both strategies to align MRI images with tissue sections stained with hypoxyprobe to demonstrate that MRI readouts align extremely well with histological staining (Figure 3). In each of the ROI used for the Nanostring spatial transcriptomic analysis (Figure 3C), we measured hypoxia in the same regions within the MRI image as a function of the *T*_1_ value, consistent with prior report of the agent.^1,124^

**Figure 1.**
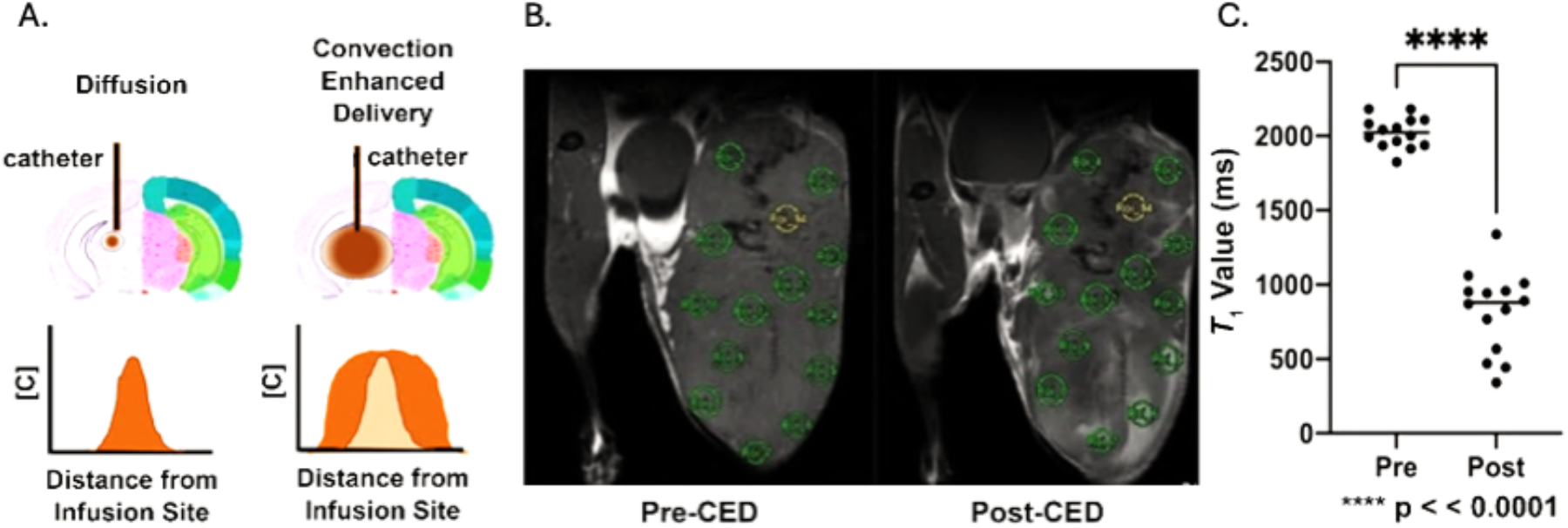
Validation of CED delivery of MRI Contrast Agents throughout the entire tumor. (a) Figure adapted from ref 163 showing how CED delivery covers more area than simple injection through a catheter that relies upon diffusion; (b) We tested CED in a tumor and measured *T*_1_ values before and after CED infusion of Magnevist, a Gd-based extracellular contrast agent. The images shown in (b) are *T*_1_-weighted images to illustrate the enhanced positive contrast from the Magnevist and also show where the ROI were selected for the *T*_1_ measurements. We picked ROI throughout the entire tumor. Before CED, *T*_1_ values were closely clustered around 2,000 ms. After CED, all ROI exhibited decreased *T*_1_ values demonstrating that through CED, we can administer contrast agent effectively throughout the entire tumor.

**Figure 2:**
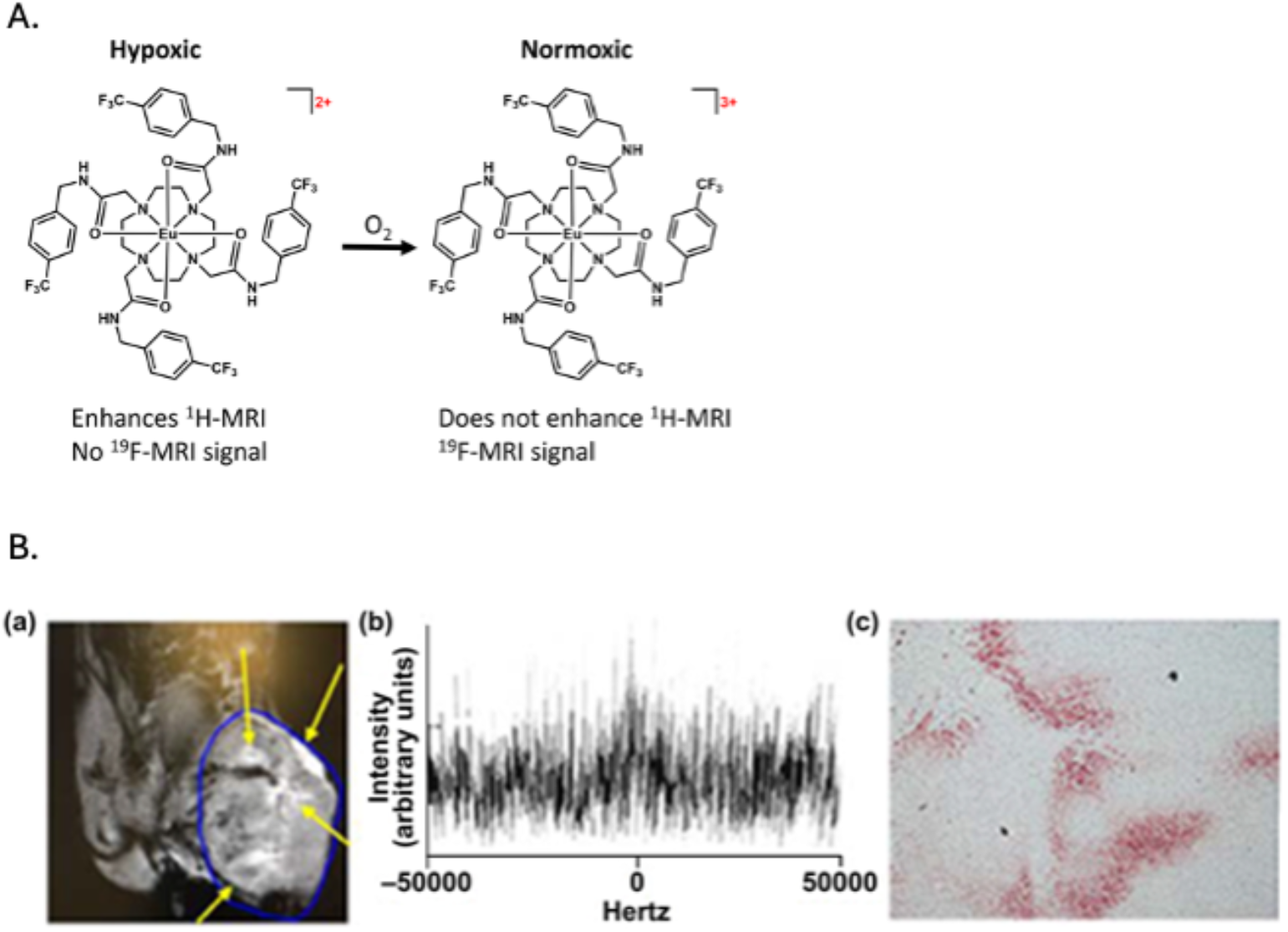
Agent mechanism and validation with hypoxyprobe staining. **A**. Schematic of how the ^19^F-Eu-based agent functions as a selective detector of hypoxia based on the oxidation state of Eu. **B**. In vivo data supporting our ability to perform the proposed studies: (a) Tumor injected with the ^19^F-Eu-based agent from **A**. Arrows identify positive *T*_1_-weighted contrast enhancement due to the agent in hypoxic regions; (b) in agreement with ^1^H-MRI, ^19^F signal is weak, indicating tumor hypoxia; and (c) hypoxyprobe histological validation of MRI data demonstrating hypoxic regions.

**Figure 3:**
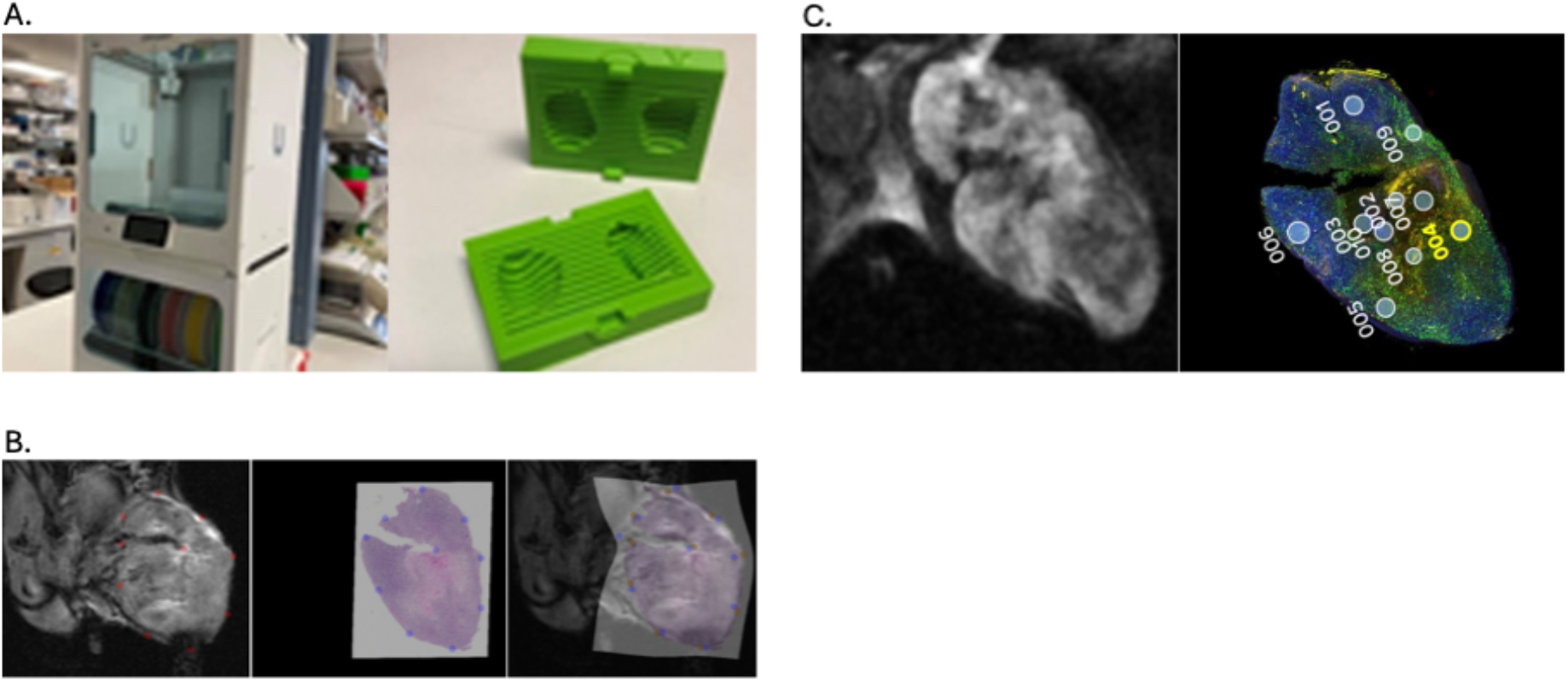
3D tumor mold for preparation and segmentation for Nanostring Analysis: A. Example of a 3D printed mold to allow for sectioning for alignment with MRI images: (Left) our high-resolution 3D printer, the Ultimaker S5 Pro; (Right) tumor mold we printed from a three-dimensional MRI dataset. We followed a reported protocol to generate the mold.^157^ **B**. (left) *T*_1_-weighted MRI image of an osteosarcoma tumor injected with our ^19^F-Eu-based agent; (center) Histological slice stained with hypoxyprobe highlighting hypoxic regions in dark pink; (right) Warping of the histological slice using RadPathFusion to match the MRI image. Note that the areas of hypoxia shown by the MRI image match well with the hypoxyprobe histology image. **C**. (left) Example MRI image of an osteosarcoma with the ^19^F-Eu-based agent infused and (right) the subsequent Nanostring data from histological sections from the same tumor. ROI used for spatial transcriptomic assessment are shown in circles. For statistical correlation data in Figure 4, we used the same ROI for spatial transcriptomic analysis, determination of hypoxia from the MRI images, and hypoxyprobe staining.

We then used statistical association and validated the hypoxia imaging data using the preliminary Nanostring spatial transcriptomics data, which robustly distinguishes between hypoxic and normoxic ROI (Figure 4A). A clear separation emerged between the two groups of ROI when transcriptomics values were plotted in t-distributed stochastic neighbor space, a commonly used nonlinear dimensionality reduction method in transcriptomics. This hypoxia–normoxia separation validates the high sensitivity of the ^19^F-Eu-based contrast agent to quantify hypoxia and of Nanostring Digital Spatial Profiler to hypoxia-induced transcriptomic changes. As an alternative and unbiased assessment of whether distinct gene sets define hypoxic and normoxic regions, we clustered the genes in the data matrix (rows representing genes and columns representing ROI). We used hierarchical clustering and measured the similarity between genes by their expression correlation over all ROI. This assessment reveals two distinct groups of co-expressing genes that characterize normoxic and hypoxic regions (Figure 4B). Additionally, differential gene expression between hypoxic and normoxic ROI confirmed significantly higher expression of known hypoxia-induced genes (hypoxia hallmark gene set)^298^ in hypoxic ROI (Figure 4C). Follow-up gene set enrichment analysis showed an enrichment of the genes co-expressing in hypoxic ROI (Figure 4B) in steroid and sugar metabolism.

**Figure 4:**
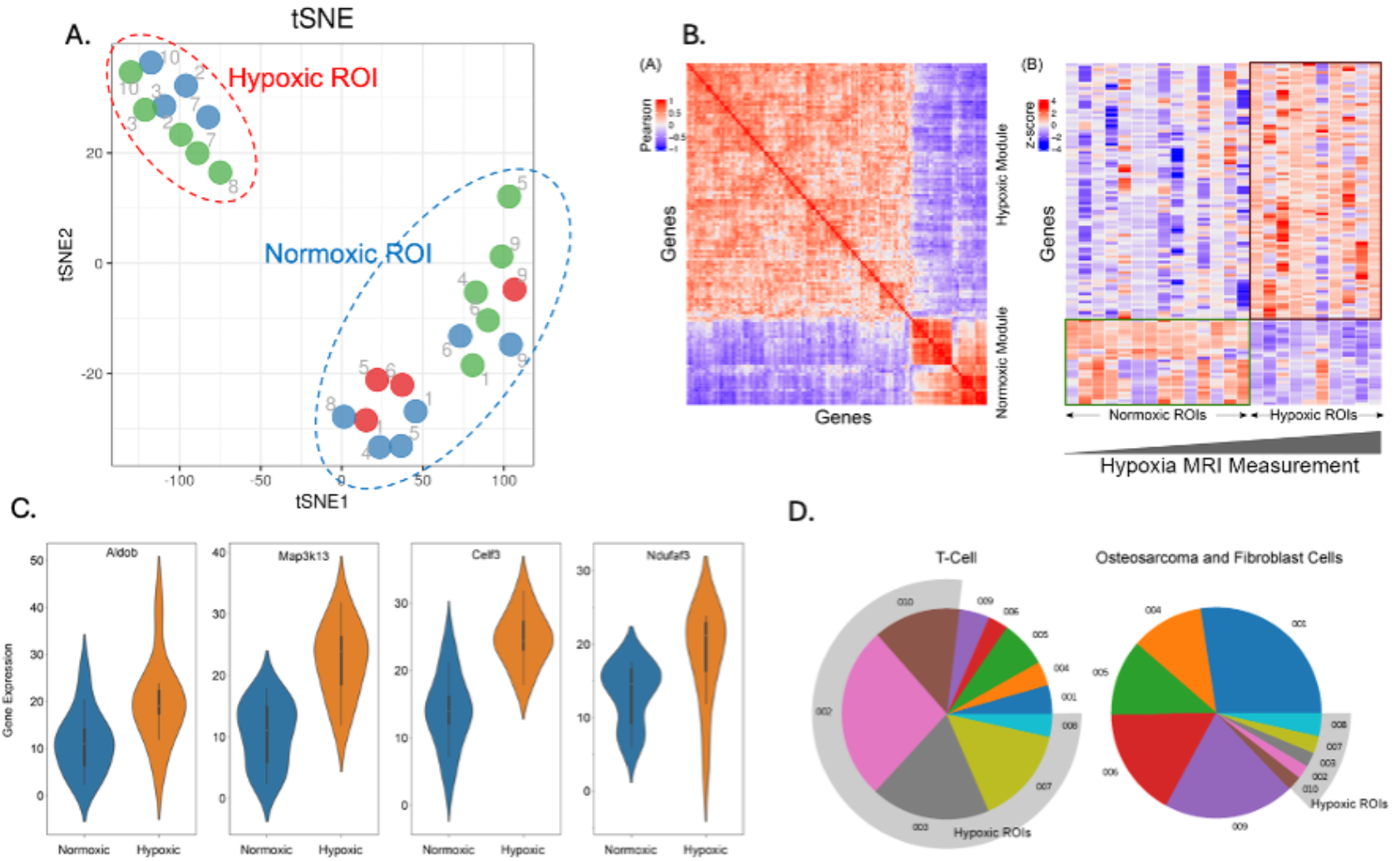
Nanostring Data: A) Nanostring data separate hypoxic and normoxic ROI. Points denote segments labeled for CD31-SMA-(green) and SMA+ (red) within ROI. B. Correlation based gene clustering: **A**. Correlation-based gene clustering (each gene is a data point, correlation of gene expression taken across ROI) revealed two groups of genes. **B**. Although the two groups were discovered without knowledge of hypoxia, they distinguish hypoxic from normoxic ROI. **C**. Differential Gene expression test between hypoxic and normoxic ROI: Differential gene expression test between hypoxic and normoxic ROI confirmed significantly high expression of known hypoxia-induced genes in hypoxic ROI (p-values < 0.05 for Aldob and Ndufaf7; < 10^−4^ for Map3k13 and Celf3). **D. Cell Type Composition Analysis:** Analysis of cell type composition reveals cell types with differential localization in hypoxic and normoxic ROI. Shown are the pie-charts of T-cell and osteosarcoma and fibroblast cell populations (different colors represent different cell types) being more localized in the hypoxic (grey outer ring) and normoxic (no outer ring) ROI, respectively.

Using a reference osteosarcoma single-cell RNA-seq (scRNA-seq) dataset,^160^ we computed cell-type proportions in all ROI. This analysis is deconvolving data of individual ROI and is routine in spatial transcriptomics studies. The hypoxic and normoxic regions show clear differences in the abundance of cell types: immune cells localize in hypoxic ROI, and fibroblasts and osteoblastic sarcoma cells localize in normoxic ROI (Figure 4D). These preliminary findings are insightful toward spatially defining cellular and molecular sarcoma landscapes.

### Discussion

In this brief research report, we demonstrate for the first time that CED is a useful tool for the distribution of both Eu-based and Gd-based contrast agents for MRI throughout a tumor. While our studies focused on intratumoral distribution, the applications of CED methods could clearly extend beyond cancer and be useful in brain tissue, muscles, and other organ systems. Although this methodology is minimally invasive, the advantages include a well-targeted administration of the agent with minimal systemic effects. An additional advantage includes a much larger distribution area of the agent compared to simple injection.

Additionally, for the first time, we also demonstrated the ability of our ^19^F-Eu-based agent to accurately and precisely identify regions of hypoxia through validation with the spatial transcriptomic Nanostring technology. Based upon the imaging readouts, we first validated that the signal generated from the ^19^F-Eu-based agent accurately distinguished hypoxic and normoxic ROI. Within these groups, we were also able to identify two distinct groups of co-expressing genes that characterized the normoxic and hypoxic regions. We observed a significantly greater levels of expression of known hypoxia-induced genes within the regions of hypoxia as well as an enrichment of steroid and sugar metabolism co-expressing genes in hypoxic ROI. Finally, we demonstrated an interesting division of cell types within normoxic and hypoxic ROI. Immune cells localize to hypoxic ROI and fibroblasts and osteoblastic sarcoma cells localized to normoxic ROI.

These results provide critical concomitant cellular and molecular insights into tumor biology directly derived from an imaging modality that can be extrapolated to the real-time radiogenomic profile of tumors. Although strategies for systemic delivery of EuII-based contrast agents are being developed,^161,162^ much work is needed to make these strategies compatible with ratiometric imaging. In the meantime, the applications of the noninvasive approach described here for the real-time detection, monitoring, and quantification of hypoxia can have a tremendous impact on patient care and management. The immediate tumor molecular landscapes identified through this novel approach can provide insights into therapeutic sensitivity at diagnosis and the evolution of tumor properties throughout the course of treatment. Thus, the integration of novel imaging strategies that relay specific tumor properties provides valuable biological insights for impacting clinical management to improve patient outcomes.

The accurate, noninvasive imaging of hypoxia is critically important because hypoxia and its consequences are attractive targets of several new drug therapies,^2–7,10–20,24–27,36,37,44,48–50^ and evaluation of the effectiveness of these therapies and their mechanism of action, even at the preclinical stage, would be enhanced with the ability to image hypoxia. Using CED with hypoxia-sensing agents such as our ^19^F-Eu-based agents hold the potential for preclinical and clinical impact.

## Acknowledgements

We would like to acknowledge the Small Animal Imaging Facility at Texas Children’s Hospital for the use of the imaging facilities. The authors gratefully acknowledge support from the National Institutes of Health (R01EB026453).

## REFERENCES

1. Basal LA, Bailey MD, Romero J, Ali MM, Kurenbekova L, Yustein J, Pautler RG, Allen MJ. Fluorinated EuII-based multimodal contrast agent for temperature- and redox-responsive magnetic resonance imaging. Chem. Sci. 2017, 8, 8345–8350.

2. Singleton DC, Macann A, Wilson WR. Therapeutic targeting of the hypoxic tumour environment. Nat. Rev. Clin. Oncol. 2021, 18, 751–772.

3. Reeves KM, Song PN, Angermeier A, Della Manna D, Li Y, Wang J, Yang ES, Sorace AG, Larimer BM. 18F-FMISO PET imaging identifies hypoxia and immunosuppressive tumor microenvironments and guides targeted Evofosfamide therapy in tumors refractory to PD-1 and CTLA-4 inhibition. Clin. Cancer Res. 2022, 28, 327–337.

4. Tao J, Yang G, Zhou W, Qiu J, Chen G, Luo W, Zhao F, You L, Zheng L, Zhang T, Zhao Y. Targeting hypoxic tumor microenvironment in pancreatic cancer. J. Hematol. Oncol. 2021, 14, 14.

5. Emami Nejad A, Najafgholian S, Rostami A, Sistani A, Shojaeifar S, Esparvarinha M, Nedaeinia R, Haghjooy Javanmard S, Taherian M, Ahmadlou M, Salehi R, Sadeghi B, Manian M. The role of hypoxia in the tumor microenvironment and development of cancer stem cell: a novel approach to developing treatment. Cancer Cell Int. 2021, 21, 62.

6. Lee P, Chandel NS, Simon MC. Cellular adaptation to hypoxia through hypoxia inducible factors and beyond. Nat. Rev. Mol. Cell Biol. 2020, 21, 268–283.

7. Wilson WR, Hay MP. Targeting hypoxia in cancer therapy. Nat. Rev. 2011, 11, 393–410.

8. Burtscher J, Mallet RT, Burtscher M, Millet GP. Hypoxia and brain aging: neurodegeneration or neuroprotection? Ageing Res. Rev. 2021, 68, 101343.

9. Lee JW, Ko J, Ju C, Eltzschig HK. Hypoxia signaling in human diseases and therapeutic targets. Exp. Mol. Med. 2019, 51, 68.

10. Chouaib S, Noman MZ, Kosmatopoulos K, Curran MA. Hypoxic stress: obstacles and opportunities for innovative immunotherapy of cancer. Oncogene 2017, 36, 439–445.

11. Bryant JL, Meredith SL, Williams KJ, White A. Targeting hypoxia in the treatment of small cell lung cancer. Lung Cancer 2014, 86, 126–132.

12. Ward C, Langdon SP, Mullen P, Harris AL, Harrison DJ, Supuran CT, Kunkler IH. New strategies for targeting the hypoxic tumour microenvironment in breast cancer. Cancer Treat. Rev. 2013, 39, 171–179.

13. Phillips RM. Targeting the hypoxic fraction of tumours using hypoxia-activated prodrugs. Cancer Chemother. Pharmacol. 2016, 77, 441–457.

14. Tatum JL. Hypoxia: importance in tumor biology, noninvasive measurement by imaging, and value of its measurement in the management of cancer therapy. Int. J. Radiat. Biol. 2006, 82, 699–757.

15. Semenza GL. Molecular mechanism mediating metastasis of hypoxic breast cancer cells. Trends Mol. Med. 2012, 18, 534–543.

16. Bredholt G, Mannelqvist M, Stefansson IM, Birkeland E, Bø TH, Øyan AM, Trovik J, Kalland KH, Jonassen I, Salvesen HB, Wik E, Akslen LA. Tumor necrosis is an important hallmark of aggressive endometrial cancer and associates with hypoxia, angiogenesis and inflammation responses. Oncotarget 2015, 6, 39676–39691.

17. Liang D, Miller GH, Tranmer GK. Hypoxia activated prodrugs: factors influencing design and development. Curr. Med. Chem. 2015, 22, 4313–4325.

18. Noman MZ, Hasmim M, Messai Y, Terry S, Kieda C, Janji B, Chouzib S. Hypoxia: A key player in antitumor immune response. A review in the theme: cellular responses to hypoxia. Am. J. Physiol.: Cell Physiol. 2015, 309, C569–C579.

19. Jain RK, Antiangiogenesis strategies revisited: from starving tumors to alleviating hypoxia. Cancer Cell 2014, 26, 605–622.

20. Harris AL. Hypoxia—a key regulatory factor in tumour growth. Nat. Rev. Cancer 2002, 2, 38–47.

21. Kunz M and Ibrahim SM. Molecular responses to hypoxia in tumor cells. Mol. Cancer 2003, 2, 23.

22. Muz B, de la Puente P, Azab F, Azab AK. The role of hypoxia in cancer progression, angiogenesis, metastasis, and resistance to therapy. Hypoxia 2015, 3, 83–92.

23. Vaupel P. Tumor microenvironmental physiology and its implications for radiation oncology. Semin. Radiat. Oncol. 2004, 14, 198–206.

24. Finger EC, Giaccia AJ. Hypoxia, inflammation, and the tumor microenvironment in metastatic disease. Cancer Metastasis Rev. 2010, 29, 285–293.

25. O’Connor LJ, Cazares-Körner C, Saha J, Evans CNG, Stratford MRL, Hammond EM, Conway SJ. Design, synthesis and evaluation of molecularly targeted hypoxia-activated prodrugs. Nat. Protoc. 2016, 11, 781–794.

26. Challapalli A, Carroll L, Aboagye EO. Molecular mechanisms of hypoxia in cancer. Clin. Transl. Imaging 2017, 5, 225–253.

27. Mehibel M, Xu Y, Li CG, Jung Moon E, Thakkar KN, Diep AN, Kim RK, Bloomstein JD, Xiao Y, Bacal J, Saldivar JC, L. Qt, Cimprich KA, Rankin EB, Giaccia AJ. Eliminating hypoxic tumor cells improves response to PARP inhibitors in homologous recombination-deficient cancer models. J. Clin. Invest. 2021, 131, e146256.

28. Bhandari V, Li CH, Bristow RG, Boutros PC, PCAWG Consortium. Divergent mutational processes distinguish hypoxic and normoxic tumours. Nat. Commun. 2020, 11, 737.

29. Jiang F, Mao Y, Lu B, Zhou G, Wang J. A hypoxia risk signature for the tumor immune microenvironment evaluation and prognosis prediction in acute myeloid leukemia. Sci. Rep. 2021, 11, 14657.

30. Schulze A, Harris AL. How cancer metabolism is tuned for proliferation and vulnerable to disruption. Nature 2012, 491, 364–373.

31. DeBerardinis RJ, Chandel NS. Fundamentals of cancer metabolism. Sci. Adv. 2016, 2, e1600200.

32. Lee DC, Sohn HA, Park ZY, Oh S, Kang YK, Lee Km, Kang M, Jang YJ, Yang SJ, Hong YK, Noh H, Kim JA, Kim DJ, Bae KH, Kim DM, Chung SJ, Yoo HS, Yu DY, Park KC, Yeom YI. A lactate-induced response to hypoxia. Cell 2015, 161, 595–609.

33. Zeng W, Liu P, Pan W, Singh SR, Wei Y. Hypoxia and hypoxia inducible factors in tumor metabolism. Cancer Lett. 2015, 356, 263–267.

34. Intlekofer AM, Dematteo RG, Venneti S, Finley LWS, Lu C, Judkins AR, Rustenburg AS, Grinaway PB, Chodera JD, Cross JR, Thompson CB. Hypoxia induces production of L-2-hydroxyglutarate. Cell Metab. 2015, 22, 304–311.

35. Thienpont B, Steinbacher J, Zhao H, D’Anna F, Kuchnio A, Ploumakis A, Ghesquière B, Van Dyck L, Boeckx B, Schoonjans L, Hermans E, Amant F, Kristensen VN, Koh KP, Mazzone M, Coleman ML, Carell T, Carmeliet P, Lambrechts D. Tumour hypoxia causes DNA hypermethylation by reducing TET activity. Nature 2016, 537, 63–68.

36. Rockwell S, Dobrucki IT, Kim EY, Marrison ST, Vu VT. Hypoxia and radiation therapy: past history, ongoing research, and future promise. Curr. Mol. Med. 2009, 9, 442–458.

37. Lu Y, Chu A, Turker MS, Glazer PM. Hypoxia-induced epigenetic regulation and silencing of the BRCA1 promoter. Mol. Cell. Biol. 2011, 31, 3339–3350.

38. Facciabene A, Peng X, Hagemann IS, Balint K, Barchetti A, Wang L, Gimotty PA, Gilks CB, Lal P, Zhang L, Coukos G. Tumour hypoxia promotes tolerance and angiogenesis via CCL28 and Treg cells. Nature 2011, 475, 226–230.

39. Chen Xs, Li Ly, Guan Yd, Yang Jm, Cheng Y. Anticancer strategies based on the metabolic profile of tumor cells: therapeutic targeting of the Warburg effect. Acta Pharmacol. Sin. 2016, 37, 1013–1019.

40. Koppenol WH, Bounds PL, Dang CV. Otto Warburg’s contributions to current concepts of cancer metabolism. Nat. Rev. Cancer 2011, 11, 325–337.

41. Vander Heiden MG, Cantley LC, Thompson CB. Understanding the Warburg effect: the metabolic requirements of cell proliferation. Science 2009, 324, 1029–1033.

42. Zhou H, Guo M, Li J, Qin F, Wang Y, Liu T, Liu J, Farhadi Sabet Z, Wang Y, Liu Y, Huo Q, Chen C. Hypoxia-triggered self-assembly of ultrasmall iron oxide nanoparticles to amplify the imaging signal of a tumor. J. Am. Chem. Soc. 2021, 143, 1846–1853.

43. Wang L, Wang H, Shen K, Park H, Zhang T, Wu X, Hu M, Yuan H, Chen Y, Wu Z, Wang Q, Li Z. Development of novel 18F-PET agents for tumor hypoxia imaging. J. Med. Chem. 2021, 64, 5593–5602.

44. Kakkad S, Krishnamachary B, Jacob D, Pacheco-Torres J, Goggins E, Kumar Bharti S, Penet MF, Bhujwalla ZM. Molecular and functional imaging insights into the role of hypoxia in cancer aggression. Cancer Metastasis Rev. 2019, 38, 51–64.

45. Brown JM. Tumor hypoxia in cancer therapy. In Oxygen Biology and Hypoxia, Methods in Enzymology Series Vol. 435, Academic Press: San Diego, 2007, Chapter 15, pp 297–321.

46. Horsman MR, Mortensen LS, Petersen JB, Busk M, Overgaard J. Imaging hypoxia to improve radiotherapy outcome. Nat. Rev. Clin. Oncol. 2012, 9, 674–687.

47. Varia MA, Calkins-Adams DP, Rinker LH, Kennedy AS, Novotny DB, Fowler WC, Raleigh JA. Pimonidazolea novel hypoxia marker for complementary study of tumor hypoxia and cell proliferation in cervical carcinoma. Gynecol. Oncol. 1998, 71, 270–277.

48. Parks SK, Chiche J, Pouysségur J. Disrupting proton dynamics and energy metabolism for cancer therapy. Nat. Rev. Cancer 2012, 13, 611–623.

49. Błaszczak-Świątkiewicz K, Mikiciuk-Olasik E. Some characteristics of activity of potential chemotherapeutics–benzimidazole derivatives. Adv. Med. Sci. 2015, 60, 125–132.

50. Sharma A, Arambula JF, Koo S, Kumar R, Singh H, Sessler JL, Seung Kim J. Hypoxia-targeted drug delivery. Chem. Soc. Rev. 2019, 48, 771–813.

51. Zheng J, Klinz SG, De Sousa R, Fitzgerald J, Jaffray DA. Longitudinal tumor hypoxia imaging with [18F]FAZA-PET provides early prediction of nanoliposomal irinotecan (nal-IRI) treatment activity. EJNMMI Res. 2015, 5, 57.

52. Vāvere AL, Lewis JS. Cu-ATSM: a radiopharmaceutical for the PET imaging of hypoxia. Dalton Trans. 2007, 4893–4902.

53. Krohn KA, Link JM, Mason RP. Molecular imaging of hypoxia. J. Nucl. Med. 2008, 49, 129S–148S.

54. Weissleder R, Schwaiger MC, Gambhir SS, Hricak H. Imaging approaches to optimize molecular therapies. Sci. Transl. Med. 2016, 8, 355ps16.

55. Mason RP, Zhao D, Pacheco-Torres J, Cui W, Kodibagkar VD, Gulaka PK, Hao G, Thorpe P, Hahn EW, Peschke P. Multimodality imaging of hypoxia in preclinical settings. Q. J. Nucl. Med. Mol. Imaging 2010, 54, 259–280.

56. Stieb S, Eleftheriou A, Warnock G, Guckenberger M, Riesterer O. Longitudinal PET imaging of tumor hypoxia during the course of radiotherapy. Eur. J. Nucl. Med. Mol. Imaging 2018, 45, 2201–2217.

57. Busk M, Overgaard J, Horsman MR. Imaging of tumor hypoxia for radiotherapy: current status and future directions. Semin. Nucl. Med. 2020, 50, 562–583.

58. Huang Y, Fan J, Li Y, Fu S, Chen Y, Wu J. Imaging of tumor hypoxia with radionuclide-labeled tracers for PET. Front. Oncol. 2021, 11, 731503.

59. Lee CT, Boss MK, Dewhirst MW. Imaging tumor hypoxia to advance radiation oncology. Antioxid. Redox Sign. 2014, 21, 313–337.

60. Walsh JC, Lebedev A, Aten E, Madsen K, Marciano L, Kolb HC. The clinical importance of assessing tumor hypoxia: relationship of tumor hypoxia to prognosis and therapeutic opportunities. Antioxid. Redox Sign. 2014, 21, 1516–1554.

61. Pacheco-Torres J, López-Larrubia P, Ballesteros P, Cerdán S. Imaging tumor hypoxia by magnetic resonance methods. NMR Biomed. 2011, 24, 1–16.

62. Gorodetskii AA, Eubank TD, Driesschaert B, Poncelet M, Ellis E, Kharmtsov VV, Bobko AA. Development of multifunctional Overhauser-enhanced magnetic resonance imaging for concurrent in vivo mapping of tumor interstitial oxygenation, acidosis and inorganic phosphate concentration. Sci. Rep. 2019, 9, 12093.

63. Chapelin F, Gedaly R, Sweeney Z, Gossett LJ. Prognostic value of fluorine-19 MRI oximetry monitoring in cancer. Mol. Imaging Biol. 2022, 24, 208–219.

64. Kodibagkar VD, Cui W, Merrritt ME, Mason RP. Novel 1H NMR approach to quantitative tissue oximetry using hexamethyldisiloxane. Magn. Reson. Med. 2006, 55, 743–748.

65. Díaz-López R, Tsapis N, Fattal E. Liquid perfluorocarbons as contrast agents for ultrasonography and 19F-MRI. Pharm. Res. 2010, 27, 1–16.

66. Evans SM, Judy KD, Dunphy I, Jenkins WT, Nelson PT, Collins R, Wileyto EP, Jenkins K, Hahn SM, Stevens CW, Judkins AR, Phillips P, Geoerger B, Koch CJ. Comparative measurements of hypoxia in human brain tumors using needle electrodes and EF5 binding. Cancer Res. 2004, 64, 1886–1892.

67. PS, Yamamoto A, Browning L, Hofer M, Adam J, Pugh CW. Recent advances in the biology of tumour hypoxia with relevance to diagnostic practice and tissue-based research. J. Pathol. 2020, 250, 593–611.

68. Gupta A, Caravan P, Price WS, Platas-Iglesias C, Gale EM. Applications for transition-metal chemistry in contrast-enhanced magnetic resonance imaging. Inorg. Chem. 2020, 59, 6648–6678.

69. Boros E, Gale EM, Caravan P. MR imaging probes: design and application. Dalton Trans. 2015, 44, 4804–4818.

70. Towner RA, Smith N, Saunders D, Henderson M, Downum K, Lupu F, Silasi-Mansat R, Ramirez DC, Gomez-Mejiba SE, Bonini MG, Ehrenshaft M, Mason RP. In vivo imaging of immuno-spin trapped radicals with molecular magnetic resonance imaging in a diabetic mouse model. Diabetes 2012, 61, 2405–2413.

71. Young SW, Qing F, Harriman A, Sessler JL, Dow WC, Mody TD, Hemmi GW, Hao Y, Miller RA. Gadolinium (III) texaphyrin: a tumor selective radiation sensitizer that is detectable by MRI. Proc. Natl. Acad. Sci. U.S.A. 1996, 93, 6610–6615.

72. Gulaka PK, Rojas-Quijano F, Kovacs Z, Mason RP, Sherry AD, Kodibagkar VD. GdDO3NI, a nitroimidazole-based T_1_ MRI contrast agent for imaging tumor hypoxia in vivo. J. Biol. Inorg. Chem. 2014, 19, 271–279.

73. Rojas-Quijano FA, Tircsó G, Benyó ET, Baranyai Z, Hoang HT, Kálmán FK, Gulaka PK, Kodibagkar VD, Aime S, Kovács Z, Sherry AD. Synthesis and characterization of a hypoxia-sensitive MRI probe. Chem.—Eur. J. 2012, 18, 9669–9676.

74. Ye D, Pandit P, Kempen P, Lin J, Xiong L, Sinclair R, Rutt B, Rao J. Redox-triggered self-assembly of gadolinium-based MRI probes for sensing reducing environment. Bioconjugate Chem. 2014, 25, 1526–1536.

75. Jagadish B, Guntle GP, Zhao D, Gokhale V, Ozumerzifon TJ, Ahad AM, Mash EA, Raghunand N. Redox-active magnetic resonance imaging contrast agents: studies with thiol-bearing 1,4,7,10-tetraazacyclododecane-1,4,7,10-tetracetic acid derivatives. J. Med. Chem. 2012, 55, 10378–10386.

76. Ye Z, Zhou Z, Ayat N, Wu X, Jin E, Shi X, Lu ZR. A neutral polydisulfide containing Gd(III) DOTA monoamide as a redox-sensitive biodegradable macromolecular MRI contrast agent. Contrast Media Mol. Imaging 2016, 11, 32–40.

77. Xie D, King TL, Banerjee A, Kohli V, Que EL. Exploiting copper redox for 19F magnetic resonance-based detection of cellular hypoxia. J. Am. Chem. Soc. 2016, 138, 2937–2940.

78. Loving GS, Mukherjee S, Caravan P. Redox-activated manganese-based MR contrast agent. J. Am. Chem. Soc. 2013, 135, 4620–4623.

79. Gale EM, Mukherjee S, Liu C, Loving GS, Caravan P. Structure-redox-relaxivity relationships for redox responsive manganese-based magnetic resonance imaging probes. Inorg. Chem. 2014, 53, 10748–10761.

80. Yu M, Beyers RJ, Gorden JD, Cross JN, Goldsmith CR. A magnetic resonance imaging contrast agent capable of detecting hydrogen peroxide. Inorg. Chem. 2012, 51, 9153–9155.

81. Tsitovich PB, Spernyak JA, Morrow JR. A redox-activated MRI contrast agent that switches between paramagnetic and diamagnetic states. Angew. Chem. Int. Ed. 2013, 52, 13997–14000.

82. Tsitovich PB, Burns PJ, McKay AM, Morrow JR. Redox-activated MRI contrast agents based on lanthanide and transition metal ions. J. Inorg. Biochem. 2014, 133, 143–154.

83. Hyodo F, Chuang KH, Goloshevshky AG, Sulima A, Griffiths GL, Mitchell JB, Koretsky AP, Krishna MC. Brain redox imaging using blood-brain barrier-permeable nitroxide MRI contrast agent. J. Celeb. Blood Flow Metab. 2008, 28, 1165–1174.

84. Do QN, Ratnakar JS, Kovács Z, Sherry AD. Redox- and hypoxia-responsive MRI contrast agents. ChemMedChem 2014, 9, 1116–1129.

85. Ratnakar SJ, Viswanathan S, Kovacs Z, Jindal AK, Green KN, Sherry AD. Europium(III) DOTA-tetraamide complexes as redox-active MRI sensors. J. Am. Chem. Soc. 2012, 134, 5798–5800.

86. Haldorsen IS, Stefansson I, Grüner R, Husby JA, Magnussen IJ, Werner HMJ, Salvesen ØO, Bjørge L, Trovik J, Taxt T, Akslen LA, Salvesen HB. Increased microvascular proliferation is negatively correlated to tumor blood flow and is associated with unfavourable outcome in endometrial carcinomas. Br. J. Cancer 2014, 110, 107–114.

87. Fusco R, Granata V, Pariante P, Cerciello V, Siani C, Di Bonito M, Valentino M, Sansone M, Botti G, Petrillo A. Blood oxygenation level dependent magnetic resonance imaging and diffusion weighted MRI imaging for benign and malignant breast cancer discrimination. Magn. Reson. Imaging 2021, 75, 51–59.

88. Dallaudiere B, Hummel V, Hess A, Lincot J, Preux PM, Maubon A, Monteil J. Tumoral hypoxia in osteosarcoma in rats: preliminary study of blood oxygenation level-dependent functional MRI and 18F-misonidazole PET/CT with diffusion-weighted MRI correlation. Am. J. Roentgenol. 2013, 200, W187–W192.

89. Huang Z, Xu X, Meng X, Hou Z, Liu F, Hua Q, Liu Q, Xiu J. Correlations between ADC values and molecular markers of Ki-67 and HIF-1α in hepatocellular carcinoma. Eur. J. Radiol. 2015, 84, 2464–2469.

90. Carmona-Bozo JC, Manavaki R, Woitek R, Torheim T, Baxter GC, Caracò C, Provenzano E, Graves MJ, Fryer TD, Patterson AJ, Gilbert FJ. Hypoxia and perfusion in breast cancer: simultaneous assessment using PET/MR imaging. Eur. Radiol. 2021, 31, 333–344.

91. Swartz JE, Driessen JP, van Kempen PMW, de Bree R, Janssen LM, Pameijer FA, Terhaard CHJ, Philippens MEP, Willems S. Influence of tumor and microenvironment characteristics on diffusion-weighted imaging in oropharyngeal carcinoma: a pilot study. Oral Oncol. 2018, 77, 9–15.

92. Hompland T, Ellingsen C, Galappathi K, Rofstad EK. DW-MRI in assessment of the hypoxic fraction, interstitial fluid pressure, and metastatic propensity of melanoma xenografts. BMC Cancer 2014, 14, 92.

93. Hino-Shishikura A, Tateishi U, Shibata H, Yoneyama T, Nishii T, Torii I, Tateishi K, Ohtake M, Kawahara N, Inoue T. Tumor hypoxia and microscopic diffusion capacity in brain tumors: a comparison of 62Cu-Diacetyl-Bis (N4-Methylthiosemicarbazone) PET/CT and diffusion-weighted MR imaging. Eur. J. Nucl. Med. Mol. Imaging 2014, 41, 1419–1427.

94. Gaustad JV, Hauge A, Wegner CS, Simonsen TG, Lund KV, Hansem LMK, Rofstad EK. DCE-MRI of tumor hypoxia and hypoxia-associated aggressiveness. Cancers 2020, 12, 1979.

95. Liu L, Hu L, Zeng Q, Peng D, Chen Z, Huang C, Liu Z, Wen Q, Zou F, Yan L. Dynamic contrast-enhanced MRI of nasopharyngeal carcinoma: correlation of quantitative dynamic contrast-enhanced magnetic resonance imaging (DCE-MRI) parameters with hypoxia-inducible factor 1α expression and tumor grade/stage. Ann. Palliat. Med. 2021, 10, 2238–2253.

96. Hillestad T, Hompland T, Fjeldbo CS, Skingen VE, Salberg UB, Aarnes EK, Nilsen A, Lund KV, Evensen TS, Kristensen GB, Stokke T, Lyng H. MRI distinguishes tumor hypoxia levels of different prognostic and biological significance in cervical cancer. Cancer Res. 2020, 80, 3993–4003.

97. Simonsen TG, Lund KV, Hompland T, Kristensen GB, Rofstad EK. DCE-MRI–derived measures of tumor hypoxia and interstitial fluid pressure predict outcomes in cervical carcinoma. Int. J. Radiat. Oncol. Biol. Phys. 2018, 102, 1193–1201.

98. Gaustad JV, Hauge A, Wegner CA, Simonsen TG, Lund KV, Hansem LMK, Rofstad EK. DCE-MRI of tumor hypoxia and hypoxia-associated aggressiveness. Cancers 2020, 12, 1979.

99. Hariharan S, Welsh CH. Shortness of breath and hypoxemia after chemotherapy with carboplatin and gemcitabine. Chest 2007, 131, 1978–1981.

100. Sears SP, Carr G, Bime C. Acute and chronic respiratory failure in cancer patients. In Oncologic Critical Care. Nates JL, Price KJ. Eds. Springer: Cham, 2020, Vol. 1, Chapter 31, pp. 445–475.

101. Young AY, Shannon VR. Acute respiratory distress syndrome in cancer patients. In Oncologic Critical Care. Nates JL, Price KJ. Eds. Springer: Cham, 2020, Vol. 1, Chapter 37, pp. 557–582.

102. Tvsvgk T, Handa A, Kumar K, Mutreja D, Subramanian S. Chemotherapy-associated pulmonary toxicity—case series from a single center. South Asian J. Cancer 2021, 10, 255–260.

103. Krzystek-Korpacka M, Matusiewicz M, Diakowska D, Grabowski K, Blachut K, Kustrzeba-Wojcicka I, Gamian A. Impact of systemic hypoxemia on cancer aggressiveness and circulating vascular endothelial growth factors A and C in gastroesophageal cancer patients with chronic respiratory insufficiency. Exp. Oncol. 2007, 29, 236–242.

104. Marinacci LX, Simeone FJ, Lundquist AL, Kuter DJ, Mahowald GK. Case 38-2020: a 52-year-old man with cancer and acute hypoxemia. N. Engl. J. Med. 2020, 383, 2372–2383.

105. Caravan P, Tóth É, Rockenbauer A, Merbach AE. Nuclear and electronic relaxation of Eu(2+)(aq): an extremely labile aqua ion. J. Am. Chem. Soc. 1999, 121, 10403–10409.

106. Tóth É, Burai L, Merbach AE. Similarities and differences between the isoelectronic GdIII and EuII complexes with regard to MRI contrast agent applications. Coord. Chem. Rev. 2001, 216–217, 363–382.

107. Garcia J, Neelavalli J, Haacke EM, Allen MJ. Eu(II)-containing cryptates as contrast agents for ultra-high field strength magnetic resonance imaging. Chem. Commun. 2011, 47, 12858–12860.

108. Garcia J, Allen MJ. Interaction of biphenyl-functionalized Eu(2+)-containing cryptate with albumin: Implication to contrast agents in magnetic resonance imaging. Inorg. Chim. Acta 2012, 393, 324–327.

109. Garcia J, Kuda-Wedagedara AN, Allen MJ. Physical properties of Eu(2+)-containing cryptates as contrast agents for ultra-high field magnetic resonance imaging. Eur. J. Inorg. Chem. 2012, 2012, 2135–2140.

110. Ekanger LA, Ali MM, Allen MJ. Oxidation-responsive Eu(2+/3+)-liposomal contrast agent for dual-mode magnetic resonance imaging. Chem. Commun. 2014, 50, 14835–14838.

111. Allen MJ. Aqueous lanthanide chemistry in asymmetric catalysis and magnetic resonance imaging. Synlett. 2016, 27, 1310–1317.

112. Ekanger LA, Mills DR, Ali MM, Polin LA, Shen Y, Haacke EM, Allen MJ. Spectroscopic characterization of the 3+ and 2+ oxidation states of europium in a macrocyclic tetraglycinate complex. Inorg. Chem. 2016, 55, 9981–9988.

113. Lenora CU, Carniato F, Shen Y, Latif Z, Haacke EM, Martin PD, Botta M, Allen MJ. Structural features of Europium(II)-containing cryptates that influence relaxivity. Chem.—Eur. J. 2017, 23, 15404–15414.

114. Corbin BA, Basal LA, White SA, Shen Y, Haacke EM, Fishbein KW, Allen MJ. Screening of ligands for redox-active europium using magnetic resonance imaging. Bioorg. Med. Chem. 2018, 26, 5274–5279.

115. Bailey MD, Jin GX, Carniato F, Botta M, Allen MJ. Rational design of high-relaxivity EuII-based contrast agents for magnetic resonance imaging of low-oxygen environments. Chem.—Eur. J. 2021, 27, 3114–3118.

116. Ekanger LA, Polin LA, Shen Y, Haacke EM, Martin PD, Allen MJ. A EuII-containing cryptate as a redox sensor in magnetic resonance imaging of living tissue. Angew. Chem. Int. Ed. 2015, 54, 14398–14401.

117. Basal LA, Yan Y, Shen Y, Haacke EM, Mehrmohammadi M, Allen MJ. Oxidation-responsive, Eu(II/III)-based, multimodal contrast agents for magnetic resonance and photoacoustic imaging. ACS Omega 2017, 2, 800–805.

118. Wang H, Clavijo Jordan V, Ramsay IA, Sojoodi M, Fuchs BC, Tanabe KK, Caravan P, Gale EM. Molecular magnetic resonance imaging using a redox-active iron complex. J. Am. Chem. Soc. 2019, 141, 5916–5925.

119. Erstad DJ, Ramsay IA, Clavijo Jordan V, Sojoodi M, Fuchs BC, Tanabe KK, Caravan P, Gale EM. Tumor contrast enhancement and whole-body elimination of the manganese-based magnetic resonance imaging contrast agent Mn-PyC3A. Invest. Radiol. 2019, 54, 697–703.

120. Aime S, Botta M, Gianolio E, Terreno E. A p(O_2_)-responsive MRI contrast agent based on the redox switch of manganese(II/III)-porphyrin complexes. Angew. Chem. 2000, 112, 763–766.

121. Gale EM, Jones CM, Ramsay I, Farrar CT, Caravan P. A Janus chelator enables biochemically responsive MRI contrast with exceptional dynamic range. J. Am. Chem. Soc. 2016, 138, 15861–15864.

122. Yu M, Ward MB, Franke A, Ambrose SL, Whaley ZL, Miller Bradford T, Gorden JD, Beyers RJ, Cattley RC, Ivanović-Burmazović I, Schwartz DD, Goldsmith CR. Adding a second quinol to a redox-responsive MRI contrast agent improves its relaxivity response to H_2_O_2_. Inorg. Chem. 2017, 56, 2812–2826.

123. Ekanger LA, Polin LA, Shen Y, Haacke EM, Allen MJ. Evaluation of Eu(II)-based positive contrast enhancement after intravenous, intraperitoneal, and subcutaneous injections. Contrast Media Mol. Imaging 2016, 11, 299–303.

124. Subasinghe SAAS, Ortiz CJ, Romero J, Ward CL, Sertage AG, Kurenbekova L, Yustein JT, Pautler RG, Allen MJ. Toward quantification of hypoxia using fluorinated Eu(II/III)-containing ratiometric probes. Proc. Natl. Acad. Sci. U.S.A. 2023, 120, e2220891120.

125. Meyer AH, Feldsien TM, Mezler M, Untucht C, Venugopalan R, Lefebvre DR. Novel Developments to Enable Treatment of CNS Diseases with Targeted Drug Delivery. Pharmaceutics 2023, 15, 1100.

126. Porath KA, Regan MS, Griffith JI, Jain S, Stopka SA, Burgenske DM, Bakken KK, Carlson BL, Decker PA, Vaubel RA, Dragojevic S, Mladek AC, Connors MA, Hu Z, He L, Kitange GJ, Gupta SK, Feldsien TM, Lefebvre DR, Agar NYR, Eckel-Passow JE, Reilly EB, Elmquist WF, Sarkaria JN. Convection enhanced delivery of EGFR targeting antibody-drug conjugates Serclutamab talirine and Depatux-M in glioblastoma patient-derived xenografts. Neurooncol Adv. 2022, 4, vdac130.

127. Vogelbaum MA, Aghi MK. Convection-enhanced delivery for the treatment of glioblastoma. Neuro Oncol. 2015, Suppl 2, ii3–ii8.

128. Marin BM, Porath KA, Jain S, Kim M, Conage-Pough JE, Oh JH, Miller CL, Talele S, Kitange GJ, Tian S, Burgenske DM, Mladek AC, Gupta SK, Decker PA, McMinn MH, Stopka SA, Regan MS, He L, Carlson BL, Bakken K, Burns TC, Parney IF, Giannini C, Agar NYR, Eckel-Passow JE, Cochran JR, Elmquist WF, Vaubel RA, White FM, Sarkaria JN. Heterogeneous delivery across the blood-brain barrier limits the efficacy of an EGFR-targeting antibody drug conjugate in glioblastoma. Neuro Oncol. 2021, 23, 2042–2053.

129. Hadaczek P, Kohutnicka M, Krauze MT, Bringas J, Pivirotto P, Cunningham J, Bankiewicz K. Convection-enhanced delivery of adeno-associated virus type 2 (AAV2) into the striatum and transport of AAV2 within monkey brain. Hum Gene Ther. 2006, 17, 291–302.

130. Johnston LC, Eberling J, Pivirotto P, Hadaczek P, Federoff HJ, Forsayeth J, Bankiewicz KS. Clinically relevant effects of convection-enhanced delivery of AAV2-GDNF on the dopaminergic nigrostriatal pathway in aged rhesus monkeys. Hum Gene Ther. 2009, 20, 497–510.

131. Foo CY, Munir N, Kumaria A, Akhtar Q, Bullock CJ, Narayanan A, Fu RZ. Medical Device Advances in the Treatment of Glioblastoma. Cancers 2022, 14, 5341.

132. Dalle Ore C, Coleman-Abadi C, Gupta N, Mueller S. Advances and Clinical Trials Update in the Treatment of Diffuse Intrinsic Pontine Gliomas. Pediatr Neurosurg. 2023, doi: 10.1159/000529099.

133. Mehkri Y, Woodford S, Pierre K, Dagra A, Hernandez J, Reza Hosseini Siyanaki M, Azab M, Lucke-Wold B. Focused Delivery of Chemotherapy to Augment Surgical Management of Brain Tumors. Curr. Oncol. 2022, 29, 8846–8861.

134. Vogelbaum MA, Brewer C, Barnett GH, Mohammadi AM, Peereboom DM, Ahluwalia MS, Gao S. First-in-humanevaluation of the Cleveland Multiport Catheter for convection-enhanced delivery of topotecan in recurrent high-grade glioma:Results of pilot trial 1. J. Neurosurg. 2018, 1, 1–10.

135. Wang JL, Barth RF, Cavaliere R, Puduvalli VK, Giglio P, Lonser RR, Elder JB. Phase I trial of intracerebral convectionenhanced delivery of carboplatin for treatment of recurrent high-grade gliomas. PLoS ONE 2020, 15, e0244383.

136. Bruce JN, Fine RL, Canoll P, Yun J, Kennedy BC, Rosenfeld SS, Sands SA, Surapaneni K, Lai R, Yanes CL, Bagiella, E, De La Paz, RL. Regression of recurrent malignant gliomas with convection-enhanced delivery of topotecan. Neurosurgery 2011, 69, 1272–1280.

137. Kicielinski KP, Chiocca EA, Yu JS, Gill GM, Coffey M, Markert JM. Phase 1 clinical trial of intratumoural reovirus infusion for the treatment of recurrent malignant gliomas in adults. Mol. Ther. 2014, 22, 1056–1062.

138. Desjardins A, Gromeier M, Herndon JE, 2nd, Beaubier N, Bolognesi DP, Friedman AH, Friedman HS, McSherry F, Muscat AM, Nair S, Peters, KB, Randazzo D. Recurrent Glioblastoma Treated with Recombinant Poliovirus. N. Engl. J. Med. 2018, 379, 150–161.

139. Van Putten EHP, Kleijn A, Van Beusechem VW, Noske D, Lamers CH, De Goede AL, Idema S, Hoefnagel D, Kloezeman JJ, Fueyo J, Lang FF, Teunissen CE, Vernhout RM, Bakker C, Gerrisen W, Curiel DT, Vulto A, Lamfers MLM, Dirven CMF. Convection Enhanced Delivery of the Oncolytic Adenovirus Delta24-RGD in Patients with Recurrent GBM: A Phase I Clinical Trial Including Correlative Studies. Clin. Cancer Res. 2022, 28, 1572–1585.

140. Nwagwu CD, Immidisetti AV, Bukanowska G, Vogelbaum MA, Carbonell A. Convection-Enhanced Delivery of a First-in-Class Anti-1 Integrin Antibody for the Treatment of High-Grade Glioma Utilizing Real-Time Imaging. Pharmaceutics 2020, 13, 40.

141. Seo YE, Bu T, Saltzman WM. Nanomaterials for convection-enhanced delivery of agents to treat brain tumors. Curr. Opin. Biomed. Eng. 2017, 4, 1–12.

142. Lonser RR, Sarntinoranont M, Morrison PF, Oldfield EH. Convection-enhanced delivery to the central nervous system. J. Neurosurg. 2015, 122, 697–706.

143. Healy AT, Vogelbaum MA. Convection-enhanced drug delivery for gliomas. Surg. Neurol. Int. 2015, 6, S59–S67.

144. Allard E, Passirani C, Benoit JP. Convection-enhanced delivery of nanocarriers for the treatment of brain tumors. Biomaterials 2009, 30, 2302–2318.

145. Arshad A, Yang B, Bienemann AS, Barua NU, Wyatt MJ, Woolley M, Johnson DE, Edler KJ, Gill SS. Convection-enhanced delivery of carboplatin PLGA nanoparticles for the treatment of glioblastoma. PLoS ONE 2015, 10, e0132266.

146. Xi G, Robinson E, Mania-Farnell B, Vanin EF, Shim KW, Takao T, Allender EV, Mayanil CS, Soares MB, Ho D, Tomita T. Convection-enhanced delivery of nanodiamond drug delivery platforms for intracranial tumor treatment. Nanomedicine 2014, 10, 381–391.

147. Heiss JD, Argersinger DP, Theodore WH, Butman JA, Sato S, Khan OI. Convection-enhanced delivery of muscimol in patients with drug-resistant epilepsy. Neurosurgery 2019, 85, E4–E15.

148. Brown CB, Jacobs S, Johnson MP, Southerland C, Threatt S. Convection-enhanced delivery in the treatment of glioblastoma. Semin. Oncol. Nurs. 2018, 34, 494–500.

149. Bander ED, Ramos AD, Wembacher-Schroeder E, Ivasyk I, Thomson R, Morgenstern PF, Souweidane MM. Repeat convection-enhanced delivery for diffuse intrinsic pontine glioma. J. Neurosurg. Pediatr. 2020, 26, 661–666.

150. Lewis O, Woolley M, Johnson D, Rosser A, Barua NU, Bienemann AS, Gill SS, Evans S. Chronic, intermittent convection-enhanced delivery devices. J. Neurosci. Methods 2016, 259, 47–56.

151. Delorey TM, Ziegler CGK, Heimberg G, Normand R, Yang Y, Segerstolpe Å, Abbondanza D, Fleming SJ, Subramanian A, Montoro DT, Jagadeesh KA, Dey KK, Sen P, Slyper M, Pita-Juárez YH, Phillips D, Biermann J, Bloom-Ackermann Z, Barkas N, Ganna A, Gomez J, Melms JC, Katsyv I, Normandin E, Naderi P, Popov YV, Raju SS, Niezen S, Tsai LTY, Siddle KJ, Sud M, Tran VM, Vellarikkal SK, Wang Y, Amir-Zilberstein L, Atri DS, Beechem J, Brook OR, Chen J, Divakar P, Dorceus P, Engreitz JM, Essene A, Fitzgerald DM, Fropf R, Gazal S, Gould J, Grzyb J, Harvey T, Hecht J, Hether T, Jané-Valbuena J, Leney-Greene M, Ma H, McCabe C, McLoughlin DE, Miller EM, Muus C, Niemi M, Padera R, Pan L, Pant D, Pe’er C, Pfiffner-Borges J, Pinto CJ, Plaisted J, Reeves J, Ross M, Rudy M, Rueckert EH, Siciliano M, Sturm A, Todres E, Waghray A, Warren S, Zhang S, Zollinger DR, Cosimi L, Gupta RM, Hacohen N, Hibshoosh H, Hide W, Price AL, Rajagopal J, Tata PR, Riedel S, Szabo G, Tickle TL, Ellinor PT, Hung D, Sabeti PC, Novak R, Rogers R, Ingber DE, Jiang ZG, Juric D, Babadi M, Farhi SL, Izar B, Stone JR, Vlachos IS, Solomon IH, Ashenberg O, Porter CBM, Li B, Shalek AK, Villani AC, Rozenblatt-Rosen O, Regev A. COVID-19 tissue atlases reveal SARS-CoV-2 pathology and cellular targets. Nature 2021, 595, 107–113.

152. Nanostring. GeoMx™ DSP Example Data Analysis Quick Start, Support Document MAN-10109-01, Seattle, WA, 2019. https://www.nanostring.com/products/geomx-digital-spatial-profiler/geomx-dsp-overview/ x(accessed March 24, 2025).

153. Pelka K, Hofree M, Chen JH, Sarkizova S, Pirl JD, Jorgji V, Bejnood A, Dionne D, Ge WH, Xu KH, Chao SX, Zollinger DR, Lieb DJ, Reeves JW, Fuhrman CA, Hoang ML, Delorey T, Nguyen LT, Waldman J, Klapholz M, Wakiro I, Cohen O, Albers J, Smillie CS, Cuoco MS, Wu J, Su MJ, Yeung J, Vijaykumar B, Magnuson AM, Asinovski N, Moll T, Goder-Reiser MN, Applebaum AS, Brais LK, DelloStritto LK, Denning SL, Phillips ST, Hill EK, Meehan JK, Frederick DT, Sharova T, Kanodia A, Todres EZ, Jané-Valbuena J, Biton M, Izar B, Lambden CD, Clancy TE, Bleday R, Melnitchouk N, Irani J, Kunitake H, Berger DL, Srivastava A, Hornick JL, Ogino S, Rotem A, Vigneau S, Johnson BE, Corcoran RB, Sharpe AH, Kuchroo VK, Ng K, Giannakis M, Nieman LT, Boland GM, Aguirre AJ, Anderson AC, Rozenblatt-Rosen O, Regev A, Hacohen N. Spatially organized multicellular immune hubs in human colorectal cancer. Cell 2021, 184, 4734–4752.

154. Bhattacharya A, Hamilton AM, Furberg H, Pietzak E, Purdue MP, Troester MA, Hoadley KA, Love MI. An approach for normalization and quality control for Nanostring RNA expression data. Briefings Bioinf. 2021, 22, bbaa163.

155. Battacharya A, Hamilton AM, Furberg H, Pietzak E, Purdue MP, Troester MA, Hoadley KA, Love MI. An approach for normalization and quality control for Nanostring RNA expression data. Brief Bioinform. 2021, 22, bbaa163.

156. Risso D, Ngai J, Speed TP, Dudoit S. Normalization of RNA-seq data using factor analysis of control genes or samples. Nat. Biotechnol. 2014, 32, 896–902.

157. Jardim-Perassi BV, Huang S, Dominguez-Viqueira W, Poleszczuk J, Budzevich MM, Abdalah MA, Pillai SR, Ruiz E, Bui MM, Zuccari DAPC, Billies RJ, Martinez GV. Multiparametric MRI and coregistered histology identify tumor habitats in breast cancer mouse models. Cancer Res. 2019, 79, 3952–3964.

158. Rusu M, Purysko AS, Verma S, Kiechle J, Gollamudi J, Ghose S, Herrmann K, Gulani V, Paspulati R, Ponsky L, Böhm M, Haynes AM, Moses D, Shnier R, Delprado W, Thompson J, Stricker P, Madabhushi A. Computational imaging reveals shape differences between normal and malignant prostates on MRI. Sci. Rep. 2017, 7, 41261.

159. Antunes J, Viswanath S, Brady JT, Crawshaw B, Ros P, Steele S, Delaney CP, Paspulati R, Willis J, Madabhushi A. Coregistration of preoperative MRI with ex vivo mesorectal pathology specimens to spatially map post-treatment changes in rectal cancer onto in vivo imaging: preliminary findings. Acad. Radiol. 2018, 25, 833–841.

160. Zhou Y, Yang D, Yang Q, Lv X, Huang W, Zhou Z, Wang Y, Zhang Z, Yuan T, Ding X, Tang L, Zhang J, Yin J, Huang Y, Yu W, Wang Y, Zhou C, Su Y, He A, Sun Y, Shen Z, Qian B, Meng W, Fei J, Yao Y, Pan X, Chen P, Hu H. Single-cell RNA landscape of intratumoral heterogeneity and immunosuppressive microenvironment in advanced osteosarcoma. Nat. Commun. 2020, 11, 6322.

161. Rashid MM, Corbin BA, Jella P, Ortiz CJ, Samee MA, Pautler RG, Allen MJ. Systemic delivery of divalent europium from ligand screening with implications to direct imaging of hypoxia. J. Am. Chem. Soc. 2022, 144, 23053–23060.

162. Lutter JC, Batchev AL, Ortiz CJ, Sertage AG, Romero J, Subasinghe SAAS, Pedersen SE, Samee MAH, Pautler RG, Allen MJ. Outersphere approach to increasing the persistence of oxygen-sensitive europium(II)-containing contrast agents for magnetic resonance imaging with perfluorocarbon nanoemulsions toward imaging of hypoxia. Adv. Healthc. Mater. 2023, e2203209.

163. Steimle JD, Canozo FJG, Park M, Kadow ZA, Samee MAH, Martin JF. Decoding the PITX2-controlled genetic network in atrial fibrillation. JCI Insight 2022, 7, e158895.

